# VirusTaxo: Taxonomic classification of virus genome using multi-class hierarchical classification by k-mer enrichment

**DOI:** 10.1101/2021.04.29.442004

**Authors:** Rajan Saha Raju, Abdullah Al Nahid, Preonath Shuvo, Rashedul Islam

## Abstract

Taxonomic classification of viruses is a multi-class hierarchical classification problem, as taxonomic ranks (e.g., order, family and genus) of viruses are hierarchically structured and have multiple classes in each rank. Classification of biological sequences which are hierarchically structured with multiple classes is challenging. Here we developed a machine learning architecture, VirusTaxo, using a multi-class hierarchical classification by k-mer enrichment. VirusTaxo classifies DNA and RNA viruses to their taxonomic ranks using genome sequence. To assign taxonomic ranks, VirusTaxo extracts k-mers from genome sequence and creates bag-of-k-mers for each class in a rank. VirusTaxo uses a top-down hierarchical classification approach and accurately assigns the order, family and genus of a virus from the genome sequence. The average accuracies of VirusTaxo for DNA viruses are 99% (order), 98% (family) and 95% (genus) and for RNA viruses 97% (order), 96% (family) and 82% (genus). VirusTaxo can be used to detect taxonomy of novel viruses using full length genome or contig sequences.

**Availability:** Online version of VirusTaxo is available at https://omics-lab.com/virustaxo/.

## Introduction

Virus genome consists of either DNA or RNA and is broadly classified as DNA virus or RNA virus (Chaitanya, 2019). Viruses are classified into taxonomic ranks which play an important role in finding out their source, genetic relationship, ancestry and origin. Taxonomic classification of viruses ensures the consistent and accurate classification of novel viruses. Using sequencing technologies, there are methods available for automating the classification of the viruses from genomic sequences (Remita *et al.*, 2017; Vilsker *et al.*, 2019). Most of the existing methods are based on similarities in genome structure and organisation, the presence of homologous gene and protein sequences (Vilsker *et al.*, 2019; Simmonds, 2015). Homology based methods require higher computational resources and might produce unreliable alignment for novel viral species (Bazinet and Cummings, 2012). Supervised machine learning methods have been widely used to classify metagenomic reads against known viral and bacterial genomes (Ounit *et al.*, 2015; Ounit and Lonardi, 2016; Shang and Sun, 2020). Machine learning techniques have also been used to assign taxonomic labels of viruses from genome sequence in CASTOR and ML-DSP (Remita *et al.*, 2017; Randhawa *et al.*, 2020, 2019). CASTOR used the features of restriction fragment length polymorphism (Remita *et al.*, 2017) and ML-DSP used Discrete Fourier Transformation, a digital signal processing technique, to encode DNA sequence (Randhawa *et al.*, 2019). K-mer (i.e., DNA words of length k) feature is considered as core to the metagenomic read classification *(Ounit et al., 2015; Ounit and Lonardi, 2016; Lorenzi et al., 2020; Breitwieser et al., 2018)*. K-mer-based classification is shown to be powerful to detect long sequences of RNAs (Kirk *et al.*, 2018). Here we used the k-mer features to develop VirusTaxo that classifies DNA and RNA viruses to their taxonomic ranks from genome sequence. VirusTaxo has two separate models for DNA and RNA viruses to classify their taxonomic ranks. An online version of VirusTaxo is available for users to predict the taxonomic rank of a given virus from genome sequence.

## Results and discussion

### Classification of virus taxonomic ranks using VirusTaxo

We trained VirusTaxo using DNA and RNA virus genomes to predict their taxonomic ranks e.g., order, family and genus. Total 2,561 DNA and 1,480 RNA virus genomes were used to train the VirusTaxo models that belong to 231 DNA and 142 RNA virus genera **(Table 1)**. For the hierarchical classification of virus taxonomic ranks, we trained multiple classifiers at each level but during prediction we utilize one classifier at each level based on previous output except root. **(Fig. 1a)**illustrates an example of total 8 classifiers trained for 2 orders, 5 families with 17 genera at three taxonomic ranks.

**Figure 1:**
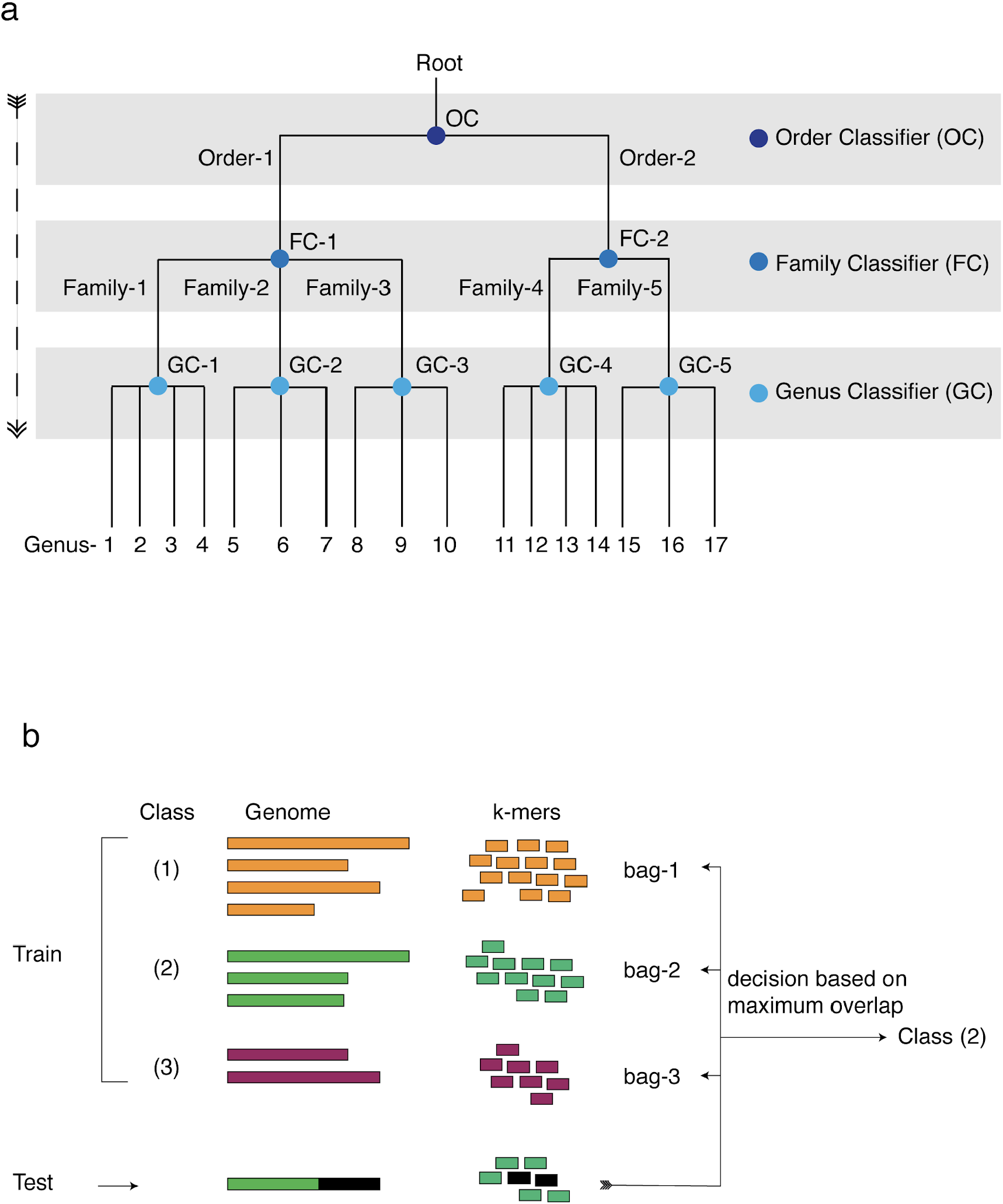
Multi-class hierarchical classification model. **a)**Example of a hierarchical structure of virus taxonomic ranks. Classifier(s) are added at each layer of taxonomic ranks. **b)**Schematic representation of the classifier for training and testing by creating bags of k-mers from virus genomes.

DNA and RNA virus genomes are different in their genome sizes (median size of DNA virus genome is 44,559bp and RNA is 8,746bp) and sequence compositions (Chaitanya, 2019). We extracted k-mers using different ranges of k-mer lengths e.g., 17-26bp and 13-22bp for DNA and RNA viruses respectively. The accuracies of DNA and RNA models varied at different k-mer lengths. K-mer lengths of 21-23bp showed highest accuracies in order (99.57%), family (98.27%) and genus (94.81%) level in the DNA model **(Fig. 2a)**. At the family level the accuracies did not change between the k-mer lengths of 20-26bp. For the RNA model, k-mer length of 17bp provided the maximum accuracies where the accuracies fluctuate with the k-mer lengths **(Fig. 2a)**. At a fixed k-mer length, the accuracies also reduced with the increase of minimum frequency threshold (MFT) of k-mers in both models **(Fig. 2b) (see methods)**. To build the prediction models, we used k-mer lengths of 21bp and 17bp for DNA and RNA virus datasets respectively with MFT value of 1. We tested the accuracies of these models to predict order, family and genus by using test datasets that contain one species genome from each genus. Total 231 DNA and 142 RNA genomes were randomly selected from each genus to generate test datasets and repeated the testing 10 times. The average accuracies were 99% (order), 98% (family) and 95% (genus) for the DNA viruses and 97% (order), 96% (family) and 82% (genus) for the RNA viruses **(Fig. 2c)**. Because of fewer branches and larger sample sizes in the top taxonomic rank, order level accuracies were highest in both models and the accuracies dropped gradually from family to genus level.

**Figure 2:**
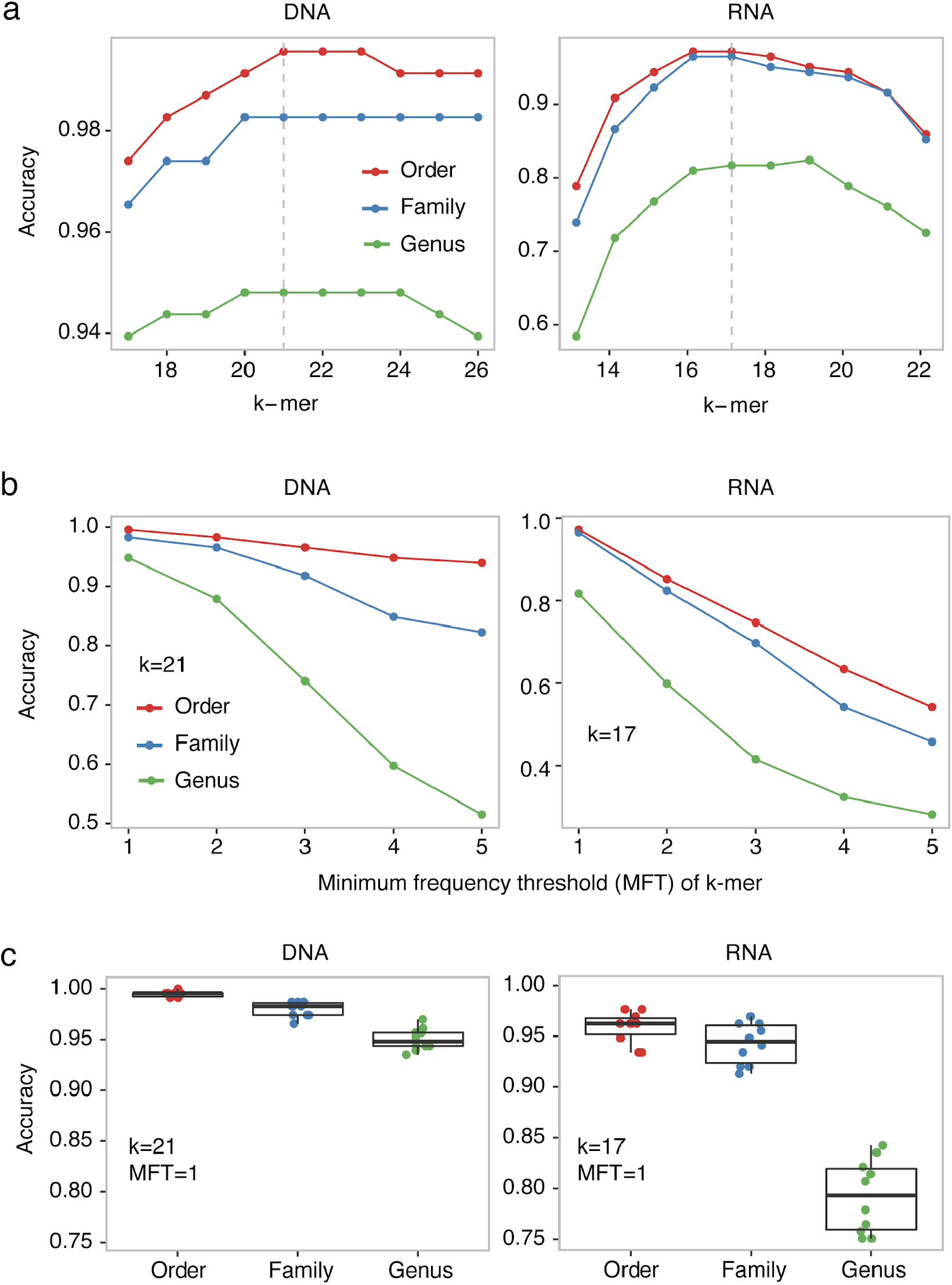
Accuracy of VirusTaxo for order, family and genus level classification. **a)**Changes of accuracies at different k-mers. For DNA and RNA datasets, 21 and 17 k-mer lengths provided the highest accuracy which is highlighted in gray dotted line. **b)**Accuracies with different minimum frequency thresholds (MFT) at k-mer length of 21bp and 17bp in DNA and RNA viruses respectively. **c)**Accuracies of VirusTaxo for 10 rounds of testing of DNA and RNA models. For each iteration of hierarchical testing, one species genome per genus was randomly selected from the DNA and RNA datasets.

### Comparison of VirusTaxo with other machine learning algorithms

To benchmark the performances of VirusTaxo with other machine learning methods, we compared VirusTaxo with four different algorithms e.g., random forest, gradient boosting, multilayer perceptron, k-nearest neighbors. In both DNA and RNA virus datasets, VirusTaxo outperformed other methods at all taxonomic ranks (e.g., order, family and genus) we analyzed. VirusTaxo showed 1% (order), 6% (family) and 16% (genus) improvement over other methods on average for DNA dataset **(Table 1)**. For RNA datasets, VirusTaxo showed 5.5% (order), 13% (family) and 27% (genus) improvement. On average RNA virus genome is 5 times smaller than DNA virus and has 43% (1480/2561) less number of species genomes available compared to DNA virus. Potentially for those reasons, accuracies of all models are relatively lower in RNA dataset across all the methods compared to DNA dataset. In RNA datasets, VirusTaxo showed significant improvement with 15% overall higher accuracies at order, family and genus levels on average compared to other methods.

**Table 1:**
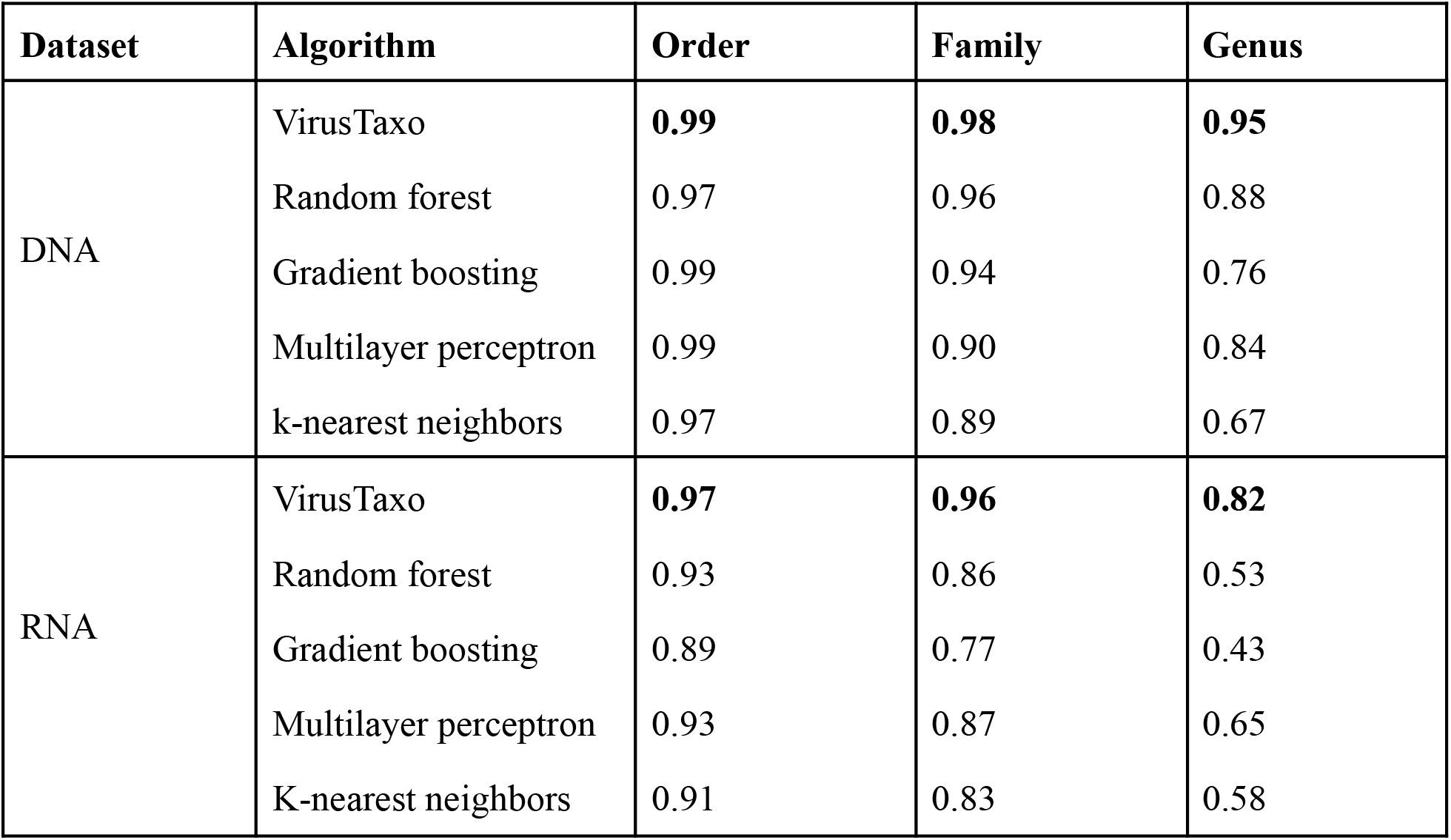
Performance comparison of different algorithms to predict virus taxonomic ranks. Performance comparison of VirusTaxo with four other algorithms using DNA and RNA datasets. Highest accuracies in each dataset are highlighted in bold font. For VirusTaxo average accuracy of 10 rounds of testing is shown here.

### Benchmarking of VirusTaxo using full and partial genome assemblies from SARS-CoV-2

SARS-CoV-2 belongs to *Betacoronavirus* genus, *Coronaviridae* family and *Nidovirales* order. **(Fig. 3a)**illustrated the taxonomic ranks of SARS-CoV-2 and its hierarchical taxonomic classification by VirusTaxo. VirusTaxo accurately identified the *Nidovirales* order, *Coronaviridae* family and *Betacoronavirus* genus from SARS-CoV-2 reference genome sequence. In addition, we obtained 5,793 de novo assemblies of SARS-CoV-2 genome *(Islam et al., 2021).* This dataset contains full and partial genome assemblies with minimum contig length of 446bp (1.49% of the 29,903bp genome) and 261 assemblies had less than 75% of the genome constructed. Despite the partial genomes provided, VirusTaxo model correctly predicted *Nidovirales* as the order, *Coronaviridae* as the family, and *Betacoronavirus* as the genus for all of the assemblies.

**Figure 3:**
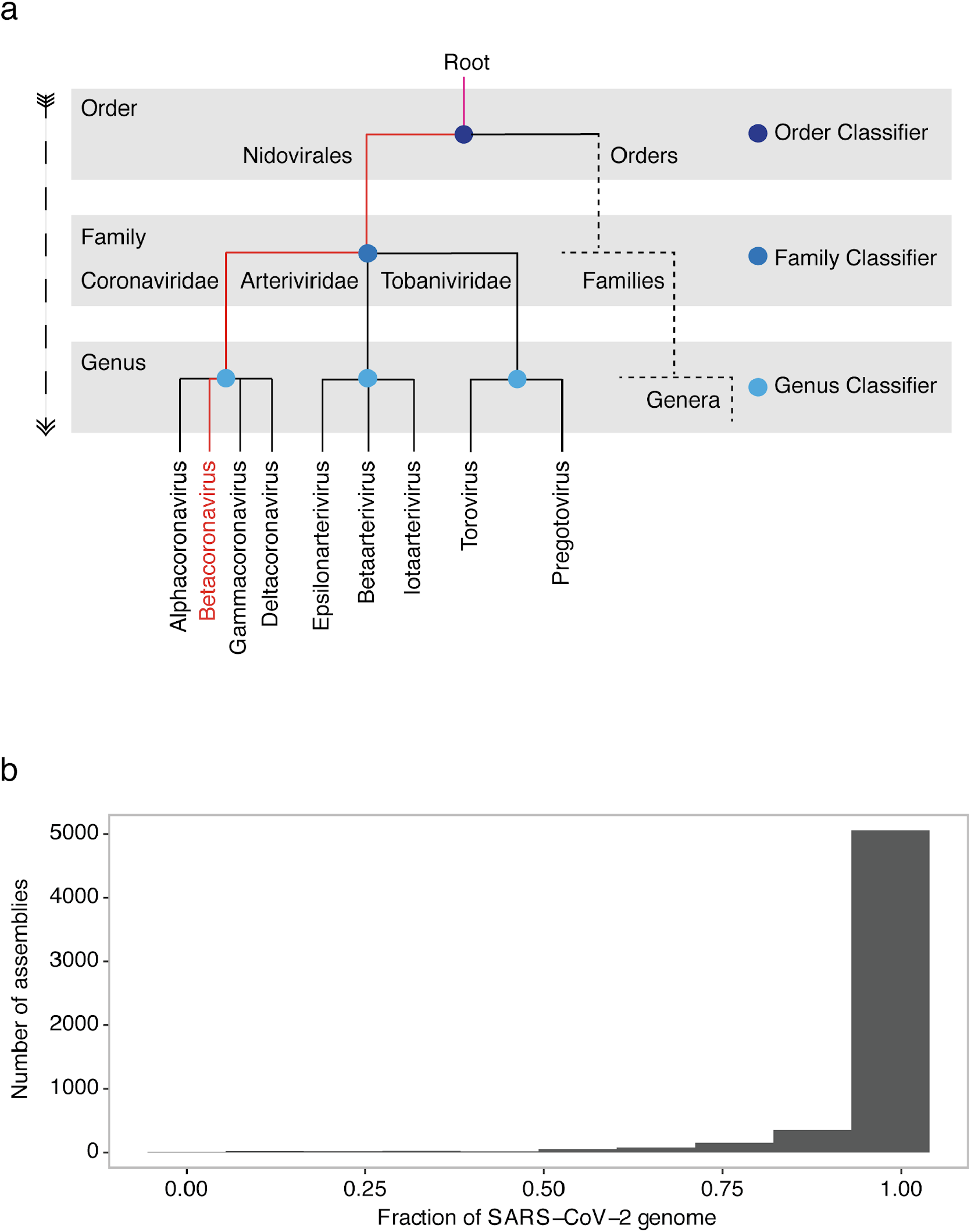
Benchmarking of VirusTaxo for SARS-CoV-2 genomes. **a)**Schematic representation of hierarchical prediction of taxonomic ranks of SARS-CoV-2 genome using VirusTaxo. In an evolutionary tree the genera were connected together by lines that come from their most recent ancestor indicating their evolutionary relationship. VirusTaxo classified the taxonomic ranks of SARS-CoV-2 for its order, family and genus which is highlighted in red color. **b)**Distribution of fraction of genome assembled in 5,793 assemblies of SARS-CoV-2 genome.

## Methods

### Datasets

RefSeq genomes of all RNA and DNA viruses were downloaded from the NCBI virus database (Brister *et al.*, 2015). Taxonomic classification of the viruses was obtained from the International Committee on Taxonomy of Viruses, ICTV Master Species List 2019.v1 release (Adams *et al.*, 2017). We chose orders with at least two families, families with at least two genera, and genera with at least three species to ensure sufficient genomes for the RNA and DNA virus classifier models. The summary of the selected datasets is listed in **(Table 1)**.

**Table 1:** Summary of RNA and DNA virus genome sequences.

|  | DNA genomes | RNA genomes |
| --- | --- | --- |
| Order | 3 | 9 |
| Family | 12 | 28 |
| Genus | 231 | 142 |
| Total species genomes | 2561 | 1480 |

### Training and testing dataset for VirusTaxo

There are 2561 and 1480 species genomes that belong to 231 and 142 unique genera of DNA and RNA viruses respectively **(Table 1)**. We randomly selected one species genome from each genus for testing the DNA and RNA models of VirusTaxo. Therefore, total 231 and 142 species genomes were used to test the DNA and RNA models respectively. For the training of DNA and RNA models, remaining 2,330 and 1,338 genomes were used in VirusTaxo. We iterated this process 10 times and the average was reported as the model accuracy.

### Hierarchical classification architecture of VirusTaxo

Each virus is associated with taxonomic ranks e.g., order, family and genus. To classify taxonomic ranks using virus genome, we developed VirusTaxo architecture of hierarchical classification of virus taxonomy illustrated in (**Fig. 1a)**. VirusTaxo uses a top to bottom approach for the classification of order, family and genus of a virus sequence. For m and n number of order and family respectively, there will be m numbers of family classifiers under respective orders in the 2nd layer and n numbers of genus classifiers under respective families in the 3rd layer. There will be a total number of classifiers = 1 + m + n in each model. We trained the classifiers at different layers of the tree by utilizing Breadth First Search (BFS) (Moore, Edward F., 1959) graph traversal algorithm for both DNA and RNA datasets. **(Fig. 1a)**illustrates an example of hierarchical classification by VirusTaxo where the order classifier (OC) in the root is classifying the genomes between two orders (e.g., Order-1 and Order-2). Then all the genomes in an order are split into corresponding families to train the family level models. Two family classifiers (e.g., FC-1 and FC-2) that belong to 2 orders classifying 5 families. Similarly, 5 genus classifiers were built to classify the genomes into 17 genera.

### Hierarchical prediction of taxonomic ranks

We pass a genome sequence through the order classifier and we get the decision for an order. Then we pass it through the family classifier under the predicted order. Finally to get the genus, we go to the genus classifier under the predicted family.

#### Training

A dataset and two parameters are required to train the multi-class classification model. These parameters are k and the minimum frequency threshold (MFT) of k-mers. Here are the training steps:

1. Create empty bags for each class.
2. Iterate over each sequence in the dataset and follow the steps mentioned below.
  a. Generate k-mers by extracting substrings of k bp with k-1 bp overlaps from the sequence.
  b. Add extracted k-mers to a bag in accordance with the class.
3. Follow the steps below for each bag.
  a. Create an empty set called temporary.
  b. Iterate over each k-mer in the bag and perform the following action.
    i. If the frequency of a k-mer in the current bag is greater than the MFT, then add it to the temporary.
  c. Update the current bag by temporary.
4. Return the bags as a model.

Training pseudocode:

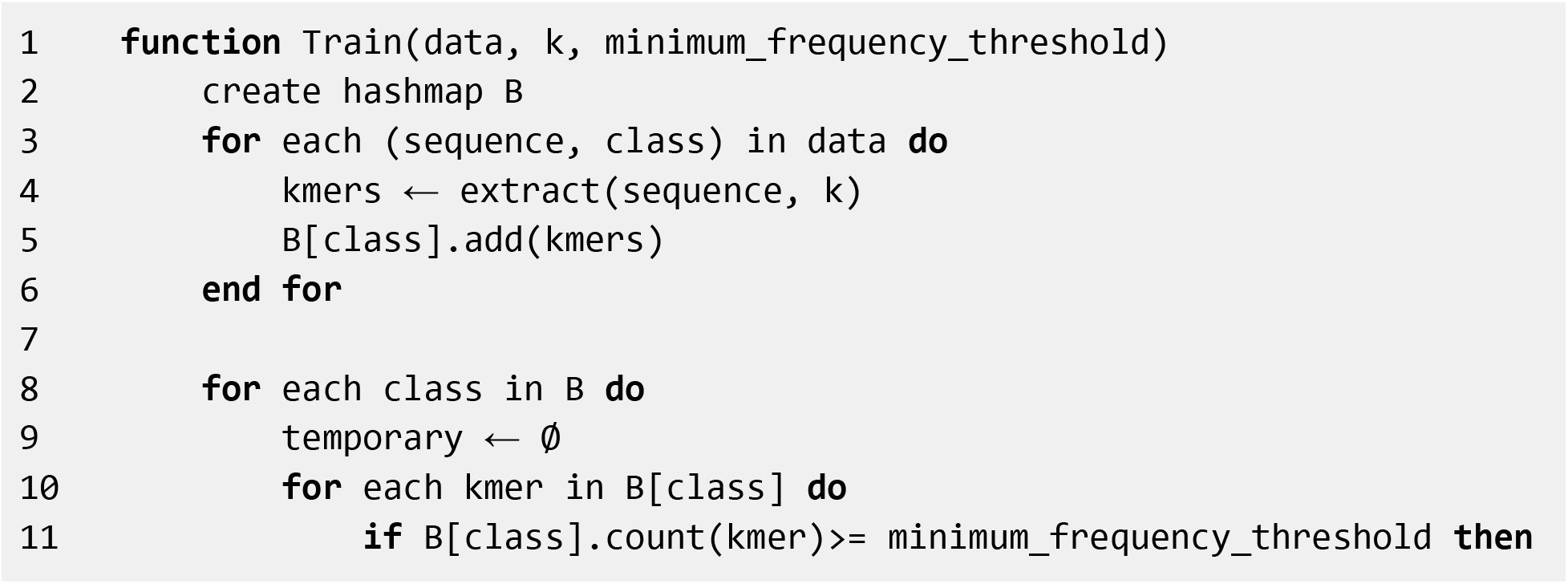
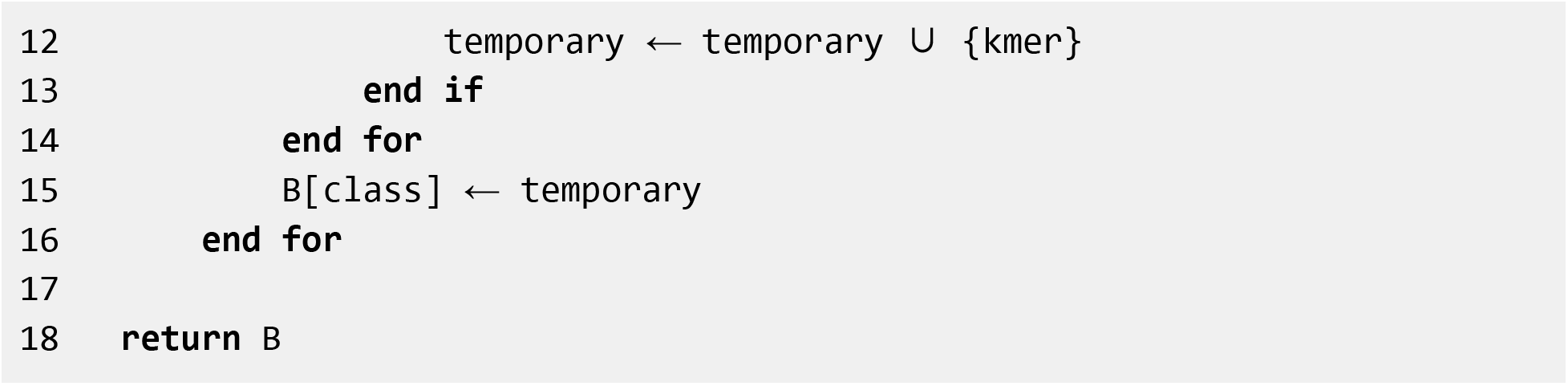

#### Prediction

Given an input sequence, model and k, the following steps are performed for predicting the rank and class.

1. Generate k-mers (same k-mer generation technique that was used in training) from the given input sequence.
2. Declare two variables called maximum count and prediction. Then initialize them with 0 and null respectively.
3. Iterate over each bag from the model and follow the steps mentioned below.
  a. Count how many extracted k-mers from the input sequence overlap with the bag.
  b. If the current count is greater than the maximum count, then update the maximum count as well as the prediction by the current count and bag’s label.
4. Return the prediction as output.

Prediction pseudocode:

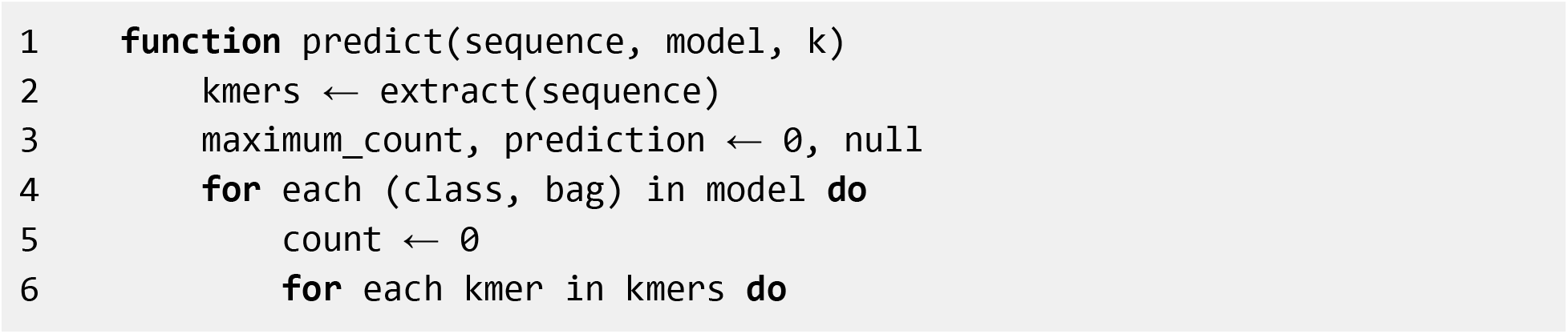
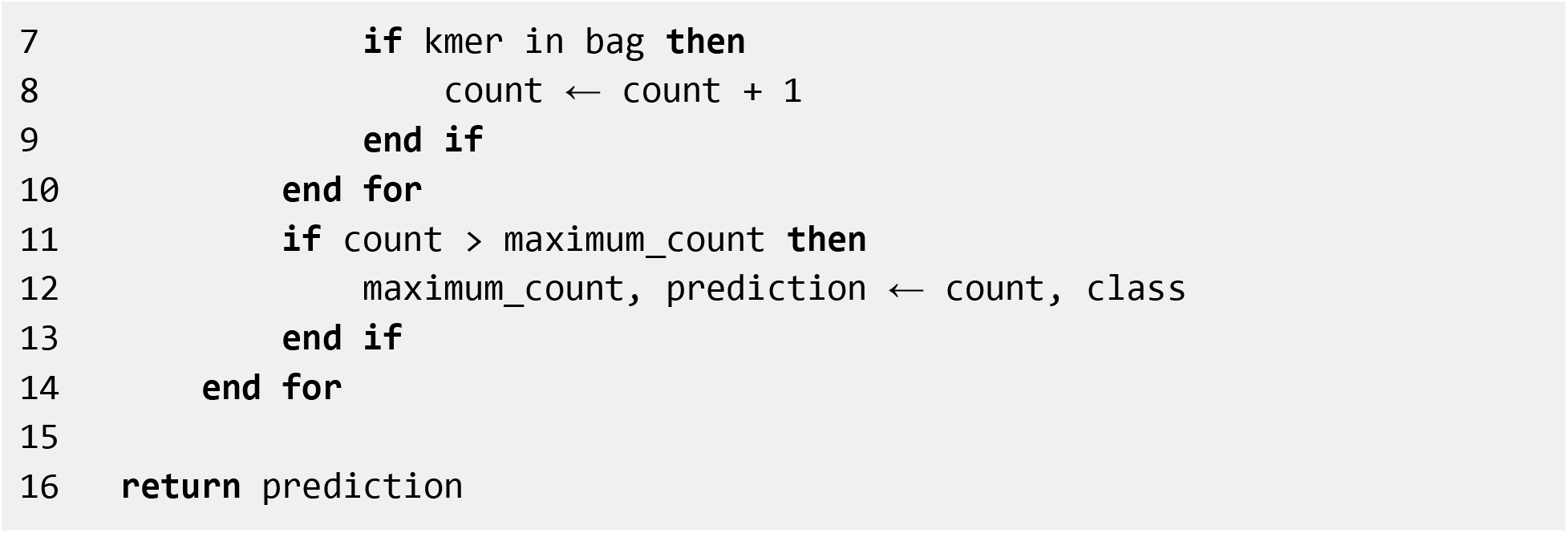

#### Benchmarking of virus taxonomic classification

We compared the performances of VirusTaxo with four other algorithms on RNA and DNA virus datasets. We used word2vec encoding (Mikolov *et al.*, 2013) for transforming genome sequences into vectors. To train two word2vec models for DNA and RNA datasets, we generated a stream of k-mers without changing sequence chronology of k-mers from each genome sequence taking 21bp and 17bp k-mer length respectively. We trained word2vec models using fastText (Bojanowski *et al.*, 2017) using the following hyperparameters in **(Table 2)**. After completion of word2vec training, we utilize four algorithms (Multilayer perceptron, Random forest, Gradient boosting, KNN) one by one in hierarchical classification of RNA virus and DNA virus. The hyperparameter details of the four algorithms are in **(Table 3)**. Here we also randomly choose one species genome from each genus to create the test set.

**Table 2:**
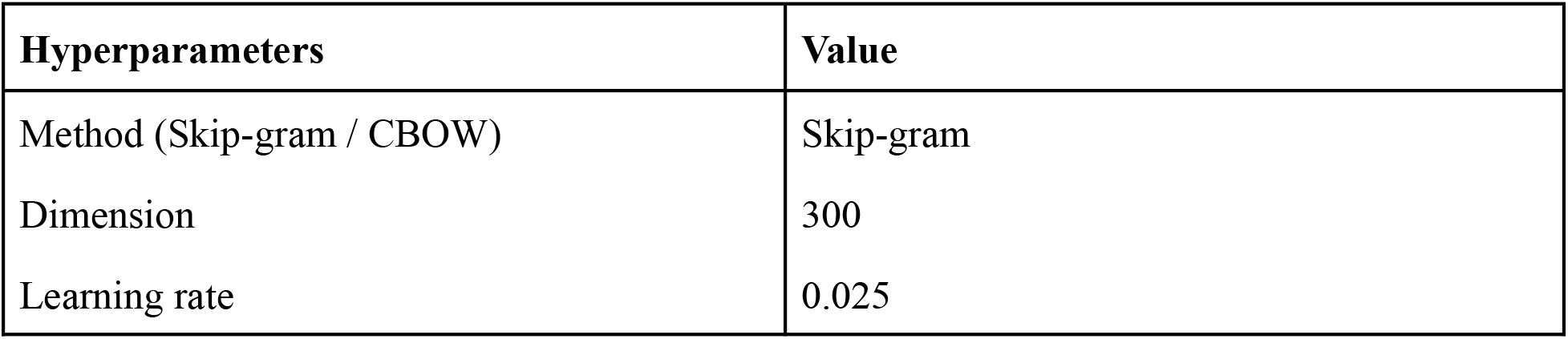
Hyperparameters setup for word2vec training.

| Hyperparameters | Value |
| --- | --- |
| Method (Skip-gram / CBOW) | Skip-gram |
| Dimension | 300 |
| Learning rate | 0.025 |

**Table 3:** Hyperparameters for four algorithms used to perform benchmarking.

| Algorithm | Hyperparameters |
| --- | --- |
| Multilayer perceptron | Hidden layer: 1<br>No of nodes in hidden layer: 100<br>Optimizer: Stochastic gradient descent<br>Learning rate: 0.01 |
| Random forest | No of trees: 100<br>Metric: Gini impurity |
| Gradient boosting | Learning rate: 0.01<br>Loss function: Deviance |
| K-nearest neighbors | K: 5 |

### Benchmarking of SARS-CoV-2 assemblies using VirusTaxo

We benchmarked the VirusTaxo RNA model using a total of 5,793 SARS-CoV-2 RNA virus assemblies from (Islam *et al.*, 2021). For each RNA fasta file, the prediction was done using the VirusTaxo RNA model.

## Acknowledgements

We acknowledge Omics-Lab (https://omics-lab.com/) for providing the research platform. We also thank Yusha Araf for his responsibilities on management and coordination of the research activity planning and execution.

## Funding

The author(s) received no financial support for the research, authorship, and/or publication of this article.

## Author Contributions

Conceptualization, R.S.R., P.S. and R.I.; Methodology, R.S.R., A.A.N. and R.I.; Software, R.S.R. and A.A.N.; Formal analysis, R.S.R., A.A.N. and R.I.; Investigation, R.S.R., A.A.N. and R.I.;

Data Curation, A.A.N., R.S.R., and R.I.; Writing – Original Draft, R.S.R., P.S. A.A.N., and R.I.; Writing – Review and Editing, R.S.R., P.S. A.A.N., and R.I.; Visualization, R.S.R. and R.I.; Supervision, R.I.

## Competing interests

The authors declare that they have no competing interests.

## Notes

### Competing Interest Statement

The authors have declared no competing interest.

